# Targeting MDM2 homodimer and heterodimer disruption with DRx-098D in TP53 wild-type and mutant cancer cells

**DOI:** 10.1101/2025.01.10.632324

**Authors:** Sean F. Cooke, Thomas A. Wright, Gillian Lappin, Elka Kyurkchieva, Yuan Yan Sin, Jiayue Ling, Alina Zorn, Bria O’Gorman, William Banyard, Chih-Jung Chang, Helen Wheadon, Danny T. Huang, George S. Baillie, Connor M. Blair

## Abstract

Novel pharmacological strategies capable of inhibiting pro-oncogenic MDM2 beyond its p53-dependent functions represents an increasingly attractive therapeutic strategy to treating solid and haematological cancers that are depdendent upon MDM2/MDMX, regardless of TP53 mutational status. Utilising a novel first-in-class cell-penetrating peptide disruptor of MDM2 homo- and heterodimerisation (DRx-098D), we demonstrate the anti-proliferative effect of blocking MDM2 dimerisation against a a panel of human cancer cell lines that are TP53 wild-type, mutant or null. DRx-098D elicits its anti-cancer activity via a differentiated mechanism vs. Idasanutlin (a Phase III clinical candidate MDM2-p53 small molecule inhibitor), inducing significantly superior growth inhibition against TP53 null HCT116 cells. Our preliminary data highlight, for the first time, the potential therapeutic utility of exploiting MDM2 dimerisation in TP53 wild-type and mutant cancers.

## Introduction

Growing recognition of the pro-oncogenic influence in which MDM2 and MDMX have in promoting cancer survival and metastasis, independent of p53, exemplifies the urgent unmet need for novel therapeutic approaches to exploiting MDM2 and MDMX activity (1-3). Current clinical candidate therapeutics capable of targeting MDM2 and MDMX (e.g., Idasanutlin, AMG-232, ALRN-6924) do so in a p53 dependent manner, specifically targeting MDM2 and/or MDMX’s ability to negatively regulate p53 and promote its degradation via the ubiquitin proteosome system (UPS) (4-5). Though this approach has proven successful in TP53 wild-type cancer, it does not translate in cancers harbouring a TP53 mutation (5). In this context, MDM2 and MDMX’s role in negatively regulating p53 expression/activity diminishes. Nonetheless, MDM2 and MDMX continue to drive tumourigenesis through a myriad p53-independent signaling pathways; an area of research that remains within its infancy (1-5).

MDM2 forms homodimers (MDM2:MDM2) and heterodimers (MDM2:MDMX) via its C-terminal RING domain. C-terminal RING domain dimerisation is essential in enabling MDM2 enzymatic E3 ligase activity (6-9). Consequently, pharmacologically targeting the de-stabilisation/disruption of C-terminal RING domain dimerisation is considered an attractive approach to inhibiting MDM2 E3 ligase activity, offering a potentially viable therapeutic strategy against MDM2:MDM2 and MDM2:MDMX dependent cancer, irrespective of TP53 mutational status (5,10).

DRx-098D-R is a novel investigative cell-penetrating peptide designed to disrupt MDM2:MDM2 and MDM2:MDMX at the dimerisation interface. In this proof-of-concept study, we outline DRx-098D-R’s ability to (i) bind both MDM2 and MDMX RING – C-terminal truncate proteins, (ii) disrupt intracellular MDM2 homodimerisation and MDM2:MDMX heterodimerisation, (iii) inhibit MDM2 E3 ligase activity,(iv)promote cancer cell death through upregulation of pro-apoptotic signaling, and (v) demonstrate preliminary therapeutic utility in the context of MDM2/MDMX expressing TP53 wild-type (WT) and mutant (MT) / null cancer. DRx-098D-R adds to the rapidly growing body of evidence supporting the therapeutic value of peptide-based disruptors of pro-oncogenic protein-protein interactions in cancer and precision medicine (11-13).

## Results and Discussion

### DRx-098D-R directly binds MDM2 and MDMX RING-C-terminal truncate proteins

To confirm target engagement, GST-fusion MDM2(428-C) or MDMX(428-C) protein was co-incubated with increasing concentrations [0.05 – 3 µM] of FITC labelled DRx-098D (DRx-098D-F) or DRx-097A (DRx-097A-F, negative control ‘knockout’ peptide) (Fig. 1B-C). Unlike DRx-097A-F, DRx-098D-F directly bound to immobilised MDM2 (Kd = 0.77 ±0.12 µM) and MDMX (Kd = 0.29 ± 0.06 µM). To determine whether binding the RING – C-terminal region of MDM2 would hinder its E3 ligase activity, DRx-098D-R (cell-permeable DRx-098D) was assessed in a cell-free *in vitro* MDM2 – p53 ubiquitination assay (Fig. 1D). DRx-098D-R, but not DRx-097A-R, induced a dose dependent inhibition of p53 ubiquitination. Thus, DRx-098D-R can inhibit MDM2’s E3 ligase activity, including its ability to ubiquitinate p53.

**Fig. 1:**
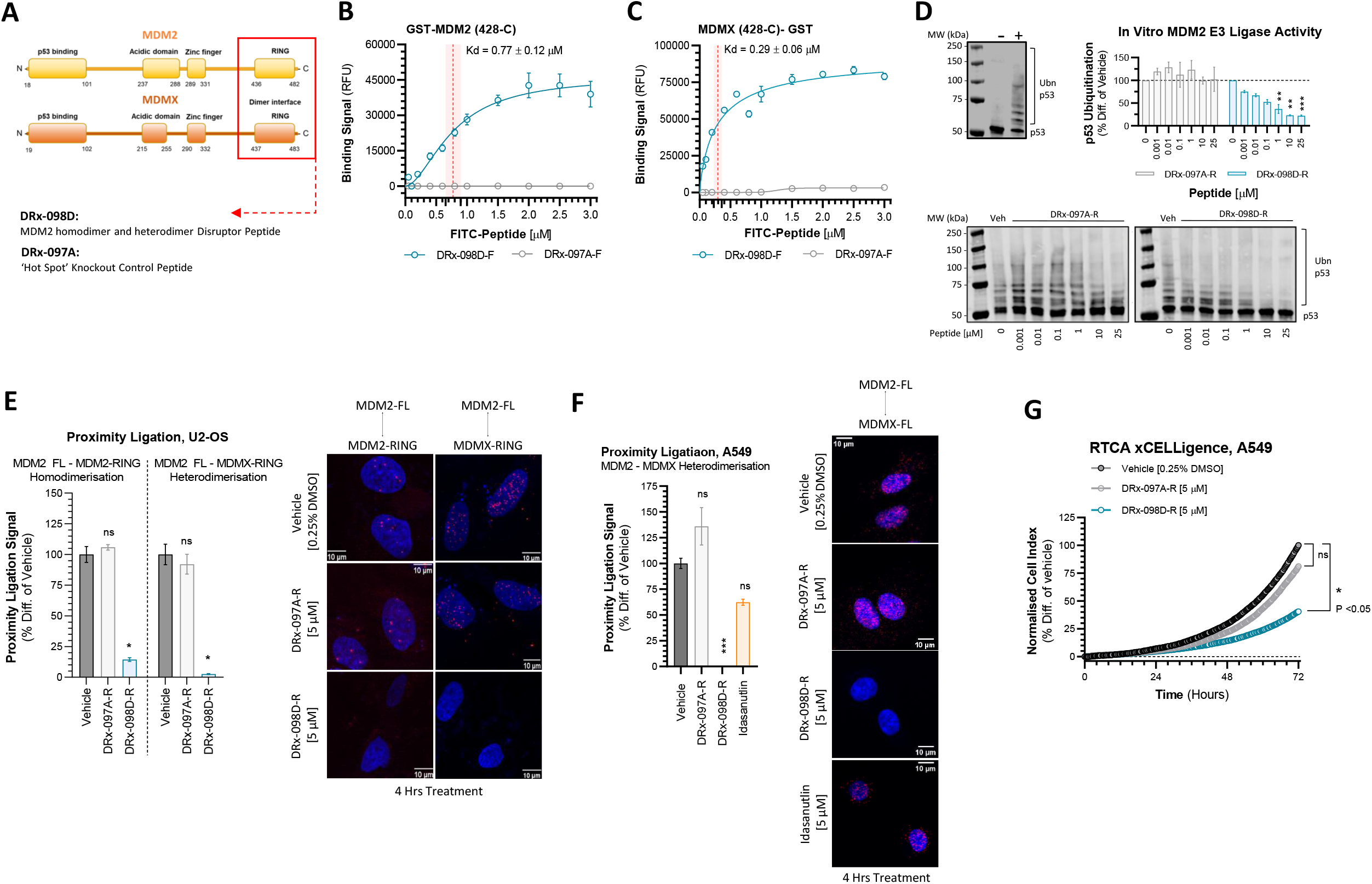
Validation of DRx-098D-R on-target mechanism. (A) MDM2 (yellow) and MDMX (orange) structural domains. Red box highlights the MDM2:MDMX RING-C-terminal domain exploited by DRx-098D-R cell-penetrating disruptor peptide. DRx-097A-R is a cell-penetrating negative peptide control containing ‘knockout’ point substitutions at key MDM2/MDMX binding residues. Target engagement assessment of (B) MDM2 (426-C)-GST and (C) MDMX (425-C)-GST with FITC labelled DRx-098D-F and DRx-097A-F. Red vertical line represents respective DRx-098D-R Kd. N=4, MEAN ± SEM. (D) Cell-free *in vitro* MDM2 E3 ligase activity assay assessing p53 ubiquitination via western immunoblotting analysis (‘+’ complete reaction mixture; ‘-’ dH_2_O in place of ATP in reaction mixture). Increasing concentrations of DRx-098D-R or DRx-097A-R were added to reaction mixture and levels of MDM2-mediated p53 ubiquitination were measured as a % difference of vehicle (DMSO). N=3, MEAN ± SEM. (E) Proximity ligation of full-length (FL) MDM2 and HA-MDM2(435-C) or Myc-MDMX(428-C) in U2-OS, treated with vehicle (0.25% DMSO), DRx-098D-R [5 µM] or DRx-097A-R [5 µM] for 4 Hrs. n=25-72, MEAN ± SEM. (F) Proximity ligation of endogenous full-length (FL) MDM2 and MDMX in A549, treated with vehicle (0.25% DMSO), DRx-098D-R [5 µM], DRx-097A-R [5 µM] or Idasanutlin [5 µM] for 4 Hrs. n=23-37, MEAN ± SEM. (G) RTCA xCELLigence analysis of A549 treated for 72 Hrs with vehicle (0.25% DMSO), DRx-098D-R [5 µM] or DRx-097A-R [5 µM]. N=3, MEAN ± SEM. *ns, not significant; *, P < 0*.*05; **, P < 0*.*01; ***, P < 0*.*001*.

### DRx-098D-R disrupts MDM2 homodimer and heterodimer formation

Given the RING – C-terminal region of MDM2 and MDMX are responsible for dimer formation, DRx-098D-R’s ability to gain intracellular access and disrupt MDM2 homodimerisation and heterodimerisation was assessed (Fig. 1E). In U2-OS cells (TP53 WT) overexpressing HA-tagged MDM2(435-C) protein, DRx-098D-R significantly downregulated complex homodimer formation with full-length endogenous MDM2. This was also observed in U2-OS cells overexpressing Myc-tagged MDMX(428-C) protein, where heterodimerisation with full-length MDM2 was significantly inhibited. Vehicle and DRx-097A-R did not affect MDM2 dimer formation (Fig. 1E). In A549 cells (TP53 WT), DRx-098D-R significantly attenuated endogenous MDM2:MDMX heterodimer formation (Fig. 1F). Again, this was not observed with vehicle or DRx-097A-R. Idasanutlin, an MDM2-p53 small molecule inhibitor, did not significantly inhibit MDM2:MDMX heterodimer formation; further differentiating DRx-098D-R’s MDM2 inhibitory mechanism. Consequently, DRx-098D-R mediated MDM2 dimer disruption significantly suppressed the relative rate of A549 cell growth vs. vehicle and DRx-097A-R (Fig. 1G). Thus, MDM2 dimer disruption is anti-proliferative in this context.

### MDM2 dimer disruption is anti-proliferative in TP53 WT and MT / null cell lines

In the context of TP53 WT cancer, current MDM2:MDMX – p53 inhibitors promote cellular apoptosis as a consequence of upregulated p53 tumour suppressor expression/activity. DRx-098D-R mediated MDM2 dimer disruption appears consistent with this mechanism, significantly upregulating MDM2, p53 and p21 protein expression (not MDMX), as well as promoting increased expression of pro-apoptotic markers Annexin V, cleaved PARP and cleaved Caspase 3 (Fig. 2A-D). However, it is well documented that targeting this pathway in TP53 MT / null cancer is ineffective, and represents a persistent barrier to existing MDM2/MDMX – p53 targeted therapeutics due to their inability to exploit MDM2/MDMX p53-independent signaling pathways.

**Fig. 2:**
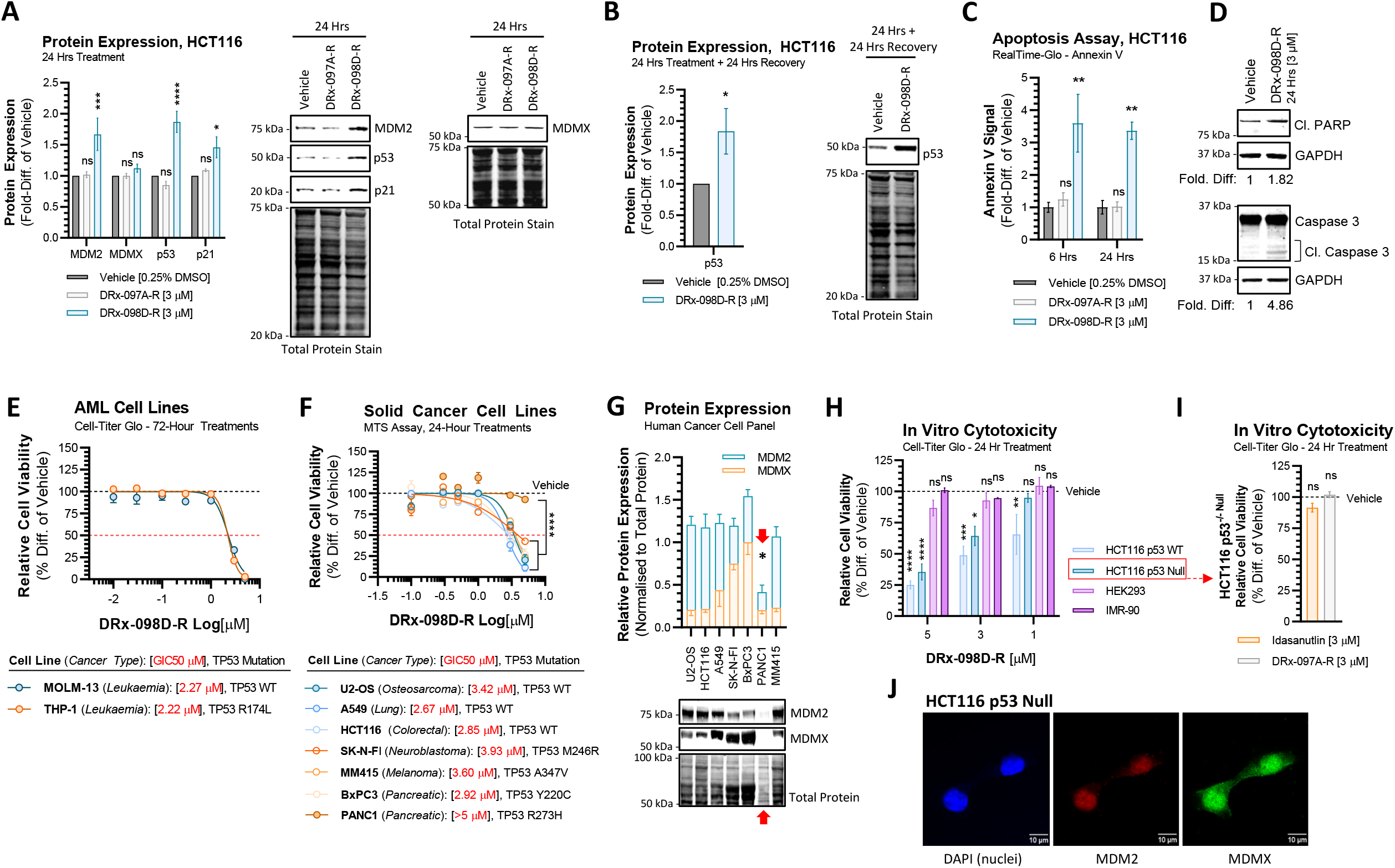
DRx-098D-R vs. TP53 wild-type and mutant human cancer cell lines. (A) Relative MDM2, MDMX, p53 and p21 protein expression in HCT116 TP53 wild-type cells following 24 Hrs treatment with vehicle (0.25% DMSO), DRx-097A-R [3 µM], DRx-098D-R [3 µM] or (B) 24 Hrs treatment followed by 24 Hrs wash-out (recovery) with vehicle (0.25% DMSO), DRx-098D-R [3 µM]. Protein expression normalised to total protein and represented as a fold difference of vehicle. N=4, MEAN ± SEM. (C) RealTime-Glo luminescence Annexin V apoptosis assay assessed relative levels of Annexin V in HCT116 TP53 wild-type cells following 6 Hrs or 24 Hrs treatment with vehicle (0.25% DMSO), DRx-097A-R [3 µM], DRx-098D-R [3 µM]. n=4, MEAN ± SEM. (D) Relative cleaved PARP and cleaved Caspase 3 protein levels in HCT116 TP53 wild-type cells following 24 Hrs treatment with vehicle (0.25% DMSO) or DRx-098D-R [3 µM]. (E) Cell-Titer Glo cell viability assessment of human acute myeloid leukaemia (AML) cell lines treated with Vehicle (0.25% DMSO) or DRx-098D-R [0.1 – 5 µM] for 72 Hrs. (F) MTS cell viability assessment of a panel of TP53 wild-type and TP53 mutant human cancer cell lines from multiple lineages, treated for 24 Hrs with vehicle (0.25% DMSO) or DRx-098D-R [0.1 – 5 µM]. Data represented as a % difference of vehicle. Black horizontal line = vehicle. Red horizontal line = growth IC50. Relative GIC50 in legend below. N=3-4, MEAN ± SEM. (G) Relative MDM2 and MDMX protein expression (untreated), normalised to total protein. Red arrow highlights MDM2/MDMX expression in PANC1 cell line. N=3-4, MEAN ± SEM. (H) Cell-Titer Glo cell viability assessment of non-cancerous HEK293 / IMR-90 and colorectal cancer HCT116 TP53 wild-type / HCT116 TP53 null cell lines follow 24 Hrs treatment with vehicle (0.25% DMSO) or DRx-098D-R [1, 3, 5 µM]. N=3-4, MEAN ± SEM. (I) Cell-Titer Glo cell viability assessment of HCT116 p53 null cells treated for 24 Hrs with vehicle (0.25% DMSO), DRx-097A-R [3 µM] or Idasanutlin [3 µM]. N=4, MEAN ± SEM. (J) HCT116 TP53 null cells immuno-fluorescently co-stained with MDM2 (red) and MDMX (green) antibodies, and counterstained with DAPI (blue). Scale bar = 10 um, n=49. *ns, not significant; *, P < 0*.*05; **, P < 0*.*01; ***, P < 0*.*001; ****, P < 0*.*0001*.

Given the growing recognition of p53-independent, pro-oncogenic MDM2/MDMX signaling, we sought to investigate the anti-proliferative consequence of disrupting MDM2 dimerisation against a panel of TP53 WT and MT human cancer cell lines from a broad range of cancer lineages (solid and haematological, Fig. 2E-F). Interestingly, DRx-098D-R significantly inhibited relative cell viability of all cell lines with similar growth IC50s, excluding PANC1 (TP53 R273W). Corresponding assessment of endogenous MDM2 and MDMX protein expression in each of the solid cancer cell lines highlighted that PANC1 expressed significantly lower levels of combined MDM2 and MDMX (Fig. 2G). These findings suggest relative combined MDM2 and MDMX protein expression may be a marker that correlates with DRx-098D-R treatment viability (i.e., DRx-098D-R activity is dependent upon the presence of MDM2/X expression)

To determine if DRx-098D-R anti-proliferative activity translated in the context of TP53 null cancer, TP53 WT and null HCT116 cancer cell lines were utilised. DRx-098D-R significantly inhibited relative cell viability of both TP53 WT and null HCT116 with similar potency (Fig. 2H). Cytotoxicity was not observed in non-cancerous HEK293 and IMR-90 cells (statistically compared to their respective vehicle control, (Fig. 2H)). As demonstrated previously with nutlin-3a, Idasanutlin had no anti-proliferative activity against HCT116 TP53 null cells (Fig. 2I) (14-15). Consistent with Fig. 2G, MDM2 and MDMX protein expression was clearly observed in HCT116 TP53 null cells (Fig. 2J).

These findings not only suggest that DRx-098D-R is inducing cancer cell-specific inhibition, but that targeted MDM2 homodimer and heterodimer disruption (i.e., by DRx-098D-R) represents a potential therapeutic vulnerability in both TP53 WT and MT cancer. This preliminary dataset warrants further development of DRx-098D-R, where particular focus should be made on its therapeutic utility in MDM2:MDM2 and MDM2:MDMX-driven TP53 MT cancer. Significant efforts should be made to characterise the potential MDM2 and/or MDMX p53-independent mechanisms at play.

## Supporting information

materials and methodology

## ACKNOWLEDGMENTS

We would like to thank Prof. Karen H. Vousden for help and advice on MDM2-MDMX-p53 cellular biology and biochemistry experiments, and for gifting plasmids and cell lines needed to carry out said experiments. This work was funded by Scottish Enterprise (PS730591C) and MRC (MC_PC_19039) grants.

## AUTHOR CONTRIBUTIONS

Conceptualization: CMB, GSB, DTH, HW. Methodology: CMB, SFC, GL, YYS, TAW. Investigation: CMB, SFC, GL, YYS, TAW, EK, AZ, JL, BG, WB, CJC. Resources: GSB, HW, DTH.

Visualization: CMB, SFC. Funding acquisition: CMB, GSB. Project Administration: CMB, SFC, GL, GSB.

Supervision: CMB, GSB. Writing – original draft: CMB, SFC. Writing – review & editing: CMB, SFC, GSB, HW, DTH.

## DATA AND MATERIALS AVAILABILITY

All relevant data needed to determine the conclusions stated within the manuscript are available in the main text or the supplementary information. Other raw data/materials used in this study are available upon request to corresponding author.

