## Supplementary material for "Targeting MDM2 homodimer and heterodimer disruption with DRx-098D in TP53 wild-type and mutant cancer cells": materials and methodology

**Supplementary Information**

**Materials and Methods:**

**Antibodies and Chemicals.** Primary antibodies include p53 (Santa Cruz, sc-126, 1:1000 – WB), MDM2 (Cell Signaling, 86934, 1:500 – WB, 1:200 – ICC/PLA), MDM2 (Abcam, ab216895, 1:50, PLA), MDMX (Sigma, HP-048821, 1:500 – WB, 1:250 – ICC/PLA), Myc (Cell Signaling, 2276, 1:200 – ICC/PLA), HA (Cell signaling, 2367, 1:200 – ICC/PLA), p21 (Santa Cruz, sc-6246, 1:1000 – WB), Caspase 3 (Cell Signaling, 9662, 1:1000 – WB), Cleaved PARP (Cell Signaling, 5625, 1:1000 – WB), GAPDH (Milipore, MAB374, 1:3000 – WB). Secondary antibodies include donkey anti-mouse 800nm (LI-COR Biosciences, 926-32212, 1:10,000 – WB), donkey anti-rabbit 800nm (LI-COR Biosciences, 926-32213, 1:10,000 – WB), goat anti-mouse 680nm (LI-COR Biosciences, 926-68070, 1:10,000 – WB), Alexa Fluor donkey anti-mouse 647nm (Thermo, A68072), Alexa Fluor donkey anti-rabbit 488nm (Thermo, A-48269). For western immunoblotting (WB), antibodies were diluted in Intercept T20 TBS antibody diluent (LI-COR). For ICC, antibodies were diluted in 5% donkey serum, 0.5% BSA in PBS. For PLA, antibodies were diluted in Duolink® antibody diluent (Merck). Stock concentrations of Idasanutlin (R&D Systems), DRx-098D (Cambridge Research Biochemicals) and DRx-097A (Cambridge Research Biochemicals) were diluted in 100% DMSO to [10 mM]. Compounds were further diluted to ≤1% DMSO in PBS or media in all assays. Unless otherwise state, drug treatments were carried out in low serum conditions (2% FBS). Idasanutlin is a clinical candidate small molecule inhibitor of the MDM2 – p53 protein-protein interaction. DRx-098D is short peptide (< 3 kDa) that has been designed to selectively target the disruption of MDM2:MDM2 homodimer and MDM2:MDMX heterodimer formation through mimicking the binding interface. DRx-098D-F is fluorescently labelled with an N-terminal FITC. DRx-098D-R possess enhanced cell permeability through addition of an optimised cationic peptide derivative at its N-terminus. DRx-097A represents the respective DRx-098D negative control peptide, where the known binding ‘hot spot’ residues have been substituted out in order to significantly negate target engagement.

**Target Engagement.** Incubated overnight at 4^o^C, 50 ng of GST-tagged MDM2(428-C) or MDMX(428-C) truncate proteins (purified as described previously (8)) were immobilised to glutathione coated wells of a pre-blocked, clear-bottom, black 96-well plate (15340, Thermo). Increasing concentrations of FITC-labelled peptide [0.05 – 3 µM] were then added to respective wells and incubated for 2 Hrs at room temperature. Wells were washed 3 times in 1X TBS-T following each incubation step to remove excess protein/peptide. Proteins and peptides were incubated in the same binding buffer (200 mM NaCl, 50 mM Tris, 5% glycerol, 5 mM DTT, 0.01% tween-20, 5 mg/mL BSA, pH 7.5). A Tristar 5 multimode microplate reader (Berthold Technologies) was utilised to measure FITC-peptide binding to MDM2 and MDMX protein. Non-linear regression analysis was performed to measure binding affinities (Kd) using GraphPad Prism 8.0 software.

C-terminal RING MDM2 protein sequence (aa428-491): SSLPLNAIEPCVICQGRPKNGCIVHGKTGHLMACFTCAKKLKKRNKPCPVCRQPIQMIVLTYFP

C-terminal RING MDMX protein sequence (aa428-490): DCQNLLKPCSLCEKRPRDGNIIHGRTGHLVTCFHCARRLKKAGASCPICKKEIQLVIKVFIA

**In Vitro Ubiquitination.** A cell-free, *in vitro*, human MDM2 ubiquitin ligase – p53 substrate kit (R&D Systems, K-200B) was used, as per manfucturer’s instructions, to assess relative MDM2 E3 ligase activity. MDM2 E3 ligase activity was assessed in the absence and presence of DRx-098D-R [0.001 – 25 µM], DRx-097A-R [0.001 – 25 µM] or Vehicle (DMSO). MDM2 E3 ligase activity was determined through p53 ubiquitination, observed/quantified utilising SDS-PAGE western immunoblotting (see protocol below). Densitometry was carried out (Image J) on non-ubiquinated p53 (i.e., single protein band at approx. 50 kDa) and the respective smear/additional bands above (i.e., ubiquinated p53). Relative p53 ubiquitination was normalised to respective non-ubiquitinated p53 and represented as a % difference of vehicle control (100%).

**Cell Culture.** PANC1 (ATCC – CRL-1469), U2-OS (ATCC – HTB-96), HCT116 TP53 wild-type, HCT116 TP53 null, SK-N-FI (ATCC – CRL-2142), IMR-90 (ATCC – CCL-186) and HEK293 (ATCC – CRL-1573) cell lines were cultured in complete DMEM. BxPC3 (ATCC – CRL-1687), MM415 (Sigma - 10092319) and A549 (ATCC – CCL-185) were cultured in complete RPMI. All media were made complete following supplementation with 2 mM L-Glutamine, 10% FBS and 100 U/mL Pen-Strep. SK-N-FI was also supplemented with 1% MEM non-essential amino acids. All cell lines were cultured in a humidified environment with 5% CO2 at 37^o^C. U2-OS, HCT116 TP53 wild-type and HCT116 TP53 null were gifted from Prof Karen H. Vousden’s research group (Francis Crick Institute, London, UK).

**Immunocytochemistry.** HCT116 p53 null cell were seeded at 0.5 x 10^5^ cells per well of a 12-well plate containing a sterilised 0.13-0.17mm glass coverslip in complete DMEM and incubated overnight. Cells were fixed in 4% paraformaldehyde (Sigma) for 15 minutes at room temperature, then permeabilised with 0.1% triton X100 (Sigma) for 4 minutes at room temperature. Cells were then blocked in 10% donkey serum, 1% BSA in PBS for 1 Hr at room temperature. Following blocking, cells were co-incubated in MDM2 (mouse) and MDMX (rabbit) primary antibodies overnight at 4^o^C. Subsequently, secondary Alexa Fluor antibodies were co-incubated for a further 1 Hr at room temperature. Cells were washed in PBS three times between each step. Following final wash, coverslips were mounted onto glass slides with Prolong Gold Antifade Mountant with DAPI (Thermo, P36941), stored in dark overnight at room temperature, and imaged using a Zeiss (LSM880) confocal microscope the following day (63X objective).

**Proximity Ligation Assay.** U2-OS cells were seeded at 1.5 x 10^5^ cells per well of a 6-well plate containing sterilised glass coverslips (0.13-0.17mm) in complete (10% FBS) media and incubated overnight. U2-OS cells were then transiently transfected with 2.5 µg (pcDNA3.1+) HA-MDM2(435-C) or Myc-MDMX(428-C) plasmid DNA (Prof. Karen H. Vousden, Francis Crick Institute, London, UK (16)) for 48 Hrs using Lipofectamine P3000 reagent as per manufacturer’s instructions (Invitrogen). U2-OS cell media was then replaced with low serum (2% FBS) media containing appropriate concentration of vehicle (0.25% DMSO), DRx-098D-R [5 µM] or DRx-097A-R [5 µM] for 4 Hrs. In contrast, A549 cells were seeded at 0.5 x 10^5^ cells per well of a 12-well plate, and cultured/treated in high serum (10% FBS) media. Following treatments, cells were fixed, permeabilised and washed as per immunocytochemistry protocol. Blocking and subsequent in situ detection of MDM2 dimerisation was then carried as per Duolink® proximity ligation manufacturer’s instructions (Merck, DUO92008). Cells were counterstained with DAPI and imaged as outlined in immunocytochemistry protocol.

C-terminal RING MDM2 sequence (aa435-491): IEPCVICQGRPKNGCIVHGKTGHLMACFTCAKKLKKRNKPCPVCRQPIQMIVLTYFP

C-terminal RING MDMX sequence (aa428-490): DCQNLLKPCSLCEKRPRDGNIIHGRTGHLVTCFHCARRLKKAGASCPICKKEIQLVIKVFIA

**RTCA xCELLigence.** Real-time cellular analysis (RTCA) of A549 cells was measured utilising the label-free cellular growth xCELLigence platform (Roche Applied Science, Agilent Technologies), where-by cellular impedance was leveraged as an indirect indicator of relative cell growth (i.e., cell index; CI). A549 cells were seeded at 1 x 10^4^ cells per well of a 96-well E-plate and allowed to adhere/grow overnight in complete (10% FBS) media. Cells were then treated with vehicle (0.25% DMSO), DRx-097A-R [5 µM] or DRx-098D-R [5 µM] for 48 Hrs, measuring cellular growth (CI) every 15 minutes. CI was normalised to 1 at treatment timepoint. Relative growth of treated A549 cells was assessed as a % difference of vehicle at 48 Hrs post-treatment (i.e., 100% relative cell growth at experiment end-point).

**Cell Viability.** For the assessment of HCT116 TP53 wild-type, HCT116 TP53 null, HEK293 and IMR-90 cells (Fig. 2A-B), and MOLM-13, THP-1 (Fig. 2E) cells, relative cell viability was assessed via CellTiter Glo 2.0 Cell Viability Assay (Promega: G9241), as per manufacturer’s instructions. For U2-OS, A549, HCT116, BxPC3, MM415, SK-N-FI, PANC1 cells (Fig. 2G), relative cell viability was assessed via CellTiter 96® Aqueous One Solution Cell Proliferation Assay (MTS, Promega: G3581), as per manufacturer’s instructions. All adherent cell lines were seeded at 5 x 10^3^ per well of a 96-well plate (MTS assay = clear plate, Cell Titer-Glo assay = clear bottom white plate) in low serum (2% FBS) media and cultured overnight. In the case of MOLM-13 and THP-1, cells were seeded at 2.5 x 10^4^ cells per well. Adherent cell lines were then treated for a further 24 Hrs in Vehicle (0.25% DMSO) or appropriate concentration of DRx-097A-R, DRx-098D-R or Idasanutlin. 0.05% triton X100 was utilised as a cytotoxic ‘kill’ control (0%). In the case of MOLM-13 and THP-1, cells were treated for a further 72 Hrs. Relative cell viability was represented as a % difference of vehicle (100%). Cell viability was measured (MTS assay = absorbance – 490nm, Cell Titer-Glo assay = luminscence) using a Tristar 5 multimode microplate reader (Berthold Technologies).

**Western Immunoblotting.** Following harvesting of protein lysates utilising appropriate lysis buffer (25 mM Tris, 150 mM NaCl, 0.1 mM EDTA, 1% NP-40, 5% glycerol, pH 7.5, protease inhibitor, phosphatase inhibitor), samples were diluted in 5X SDS gel loading buffer (10% SDS, 300 mM Tris-HCl, 0.05% bromophenol blue, 10% β-mercaptoethanol) and boiled for 5 minutes at 90^o^C. Protein resolution was carried out via SDS-PAGE (30 µg per lane, 4-12% 10-well Bis-Tris gels (NuPAGE)), transferred onto nitrocellulose membranes (GE Healthcare), and blocked in Intercept TBS blocking buffer (LI-COR). Blocked membranes were incubated in appropriate primary antibody overnight at 4^o^C, followed by 1 Hr incubation in appropriate secondary antbody at room temperature. An Odyssey CLx imaging system (LI-COR) was utilised to visualise immunoreactive bands, following which densitometry of immunoreactive bands was carried (Image J). All proteins were normalised to respective Revert^TM^ 700nm Total Protein Stain (LI-COR, 926-11011) or GAPDH.

**Annexin V Assay.** HCT116 TP53 wild-type cells were seeded at 5 x 10^3^ per well of a white, clear-bottom 96-well plate and incubated overnight. Cells were then treated with DRx-097A-R [3 µM], DRx-098D-R [3 µM] or Vehicle (0.25% DMSO) for 6 Hrs or 24 Hrs. Following treatments, Annexin V levels were measured utilising the RealTime-Glo^TM^ Annexin V Apoptosis luminescence-based assay (Promega, JA1000) as per manufacturer’s instructions. Luminescent Annexin V levels were measured via a Tristar 5 multimode microplate reader (Berthold Technologies). HCT116 cells were incubated and treated in low serum (2% FBS) conditions.

**Statistical Analysis.** All data were analysed via a one or two-way ANOVA (Dunnett’s or Tukey’s multiple comparison). Where data is represented as MEAN ± SEM from ≥3 replicates, significance was determined by a p value < 0.05. GraphPad Prism 8.0 software was utilised to statistically analysis all data.
